# Prefrontal control of visual distraction

**DOI:** 10.1101/212720

**Authors:** Joshua D. Cosman, Geoffrey F. Woodman, Jeffrey D. Schall

## Abstract

Avoiding distraction by salient irrelevant stimuli is critical to accomplishing daily tasks. Regions of prefrontal cortex control attention by enhancing the representation of task-relevant information in sensory cortex, which can be measured directly in modulation of both single neurons and averaging of the scalp-recorded electroencephalogram [1,2]. However, when irrelevant information is particularly conspicuous, it may distract attention and interfere with the selection of behaviorally relevant information. Many studies have shown that that distraction can be minimized via top-down control [3–5], but the cognitive and neural mechanisms giving rise to this control over distraction remain uncertain and vigorously debated [6–8]. Bridging neurophysiology to electrophysiology, we simultaneously recorded neurons in prefrontal cortex and event-related potentials (ERPs) over extrastriate visual cortex to track the processing of salient distractors during a visual search task. Critically, we observed robust suppression of salient distractor representations in both cortical areas, with suppression arising in prefrontal cortex before being manifest in the ERP signal over extrastriate cortex. Furthermore, only prefrontal neurons that participated in selecting the task-relevant target also showed suppression of the task-irrelevant distractor. This suggests a common prefrontal mechanism for target selection and distractor suppression, with input from prefrontal cortex being responsible for both selecting task-relevant and suppressing task-irrelevant information in sensory cortex. Taken together, our results resolve a long-standing debate over the mechanisms that prevent distraction, and provide the first evidence directly linking suppressed neural firing in prefrontal cortex with surface ERP measures of distractor suppression.

## Results

Neurons in prefrontal cortex show attention-related enhancements in firing rates to visual targets that precedes similar enhancements in extrastriate visual areas and temporal cortex [9,10]. Furthermore, causal manipulations of prefrontal cortex recapitulate this attention effect [11,12]. This suggests that input from prefrontal cortex provides an attentional control signal that gates visual processing in early sensory areas, enabling the selection of information that is relevant in a given context. However, there is a long-standing debate regarding how distracting, task-irrelevant information is processed within this system. On the one hand, stimulus-driven hypotheses propose that salient distractors automatically ‘capture’ attention and prefrontal control signals then re-direct attention to task-relevant items [6]. On the other hand, signal suppression hypotheses propose that prefrontal control signals proactively suppress the representation of salient distractors before they capture attention and interfere with the selection of task-relevant information [13,14].

This debate persists because the measures typically used to study distraction come from human behavioral and noninvasive electrophysiology studies that lack the sensitivity and specificity to resolve the dynamics of distraction control in neural systems. For example, much of this debate has played out in tasks in which observers show little or no behavioral evidence of distraction (i.e., when distraction control is effective [3–5,7,8]). Bypassing the ambiguities of behavioral evidence, electrophysiological work has sought to characterize covert responses to task-irrelevant distractors during visual search by measuring event-related potential (ERP) components putatively related to either attentional selection (the *N2pc*) or suppression (the *Pd*). However, these studies have produced mixed results, with some conditions supporting the stimulus-driven hypothesis and some supporting the signal suppression hypothesis [15–18]. One reason for these conflicting results is that the noninvasive ERP signals arise from the activity of large-scale neuronal ensembles, so the signatures of processes such as selection and suppression might overlap and mask one another. Given that top-down input from prefrontal cortex modulates processing in the extrastriate regions thought to generate the *N2pc* and *Pd* components, pairing prefrontal single unit recordings with extrastriate ERPs can resolve conflicting views of distraction control by directly measuring neuronal responses to distracting information.

To this end, we had monkeys (*Macaca radiata*) perform a visual search task in the presence or absence of a salient color distractor, while simultaneously recording neuronal discharges in frontal eye field (FEF) and event-related potentials (ERPs) over extrastriate cortex. This allowed us to track responses to both task-relevant target items and task irrelevant distractors across areas and measures in real time during visual search. Monkeys were trained to search for a T or L target in the presence or absence of a salient distractor (Figure 1), until they no longer showed a reliable influence of the distractor on either saccadic accuracy or reaction time in the previous week of training. This occurred following approximately 20 experimental sessions (~6 weeks), and mirrors the learned control over distraction observed in human studies of distraction control using an identical task [19,20]. At this point, monkeys were implanted with recording chambers and surface EEG electrodes and we began neurophysiological recordings, and across the 16 sessions reported here we found no significant effect of the salient distractor on either saccadic latency (Distractor Present = 206ms, Distractor Absent = 207ms, *t(15) < 1*) or saccadic accuracy (Distractor Present = 82.4%, Distractor Absent = 81.8%, *t(15) < 1*), indicating that the monkeys continued to exercise effective control over visual distraction during collection of the neurophysiological data reported here.

**Figure 1.**
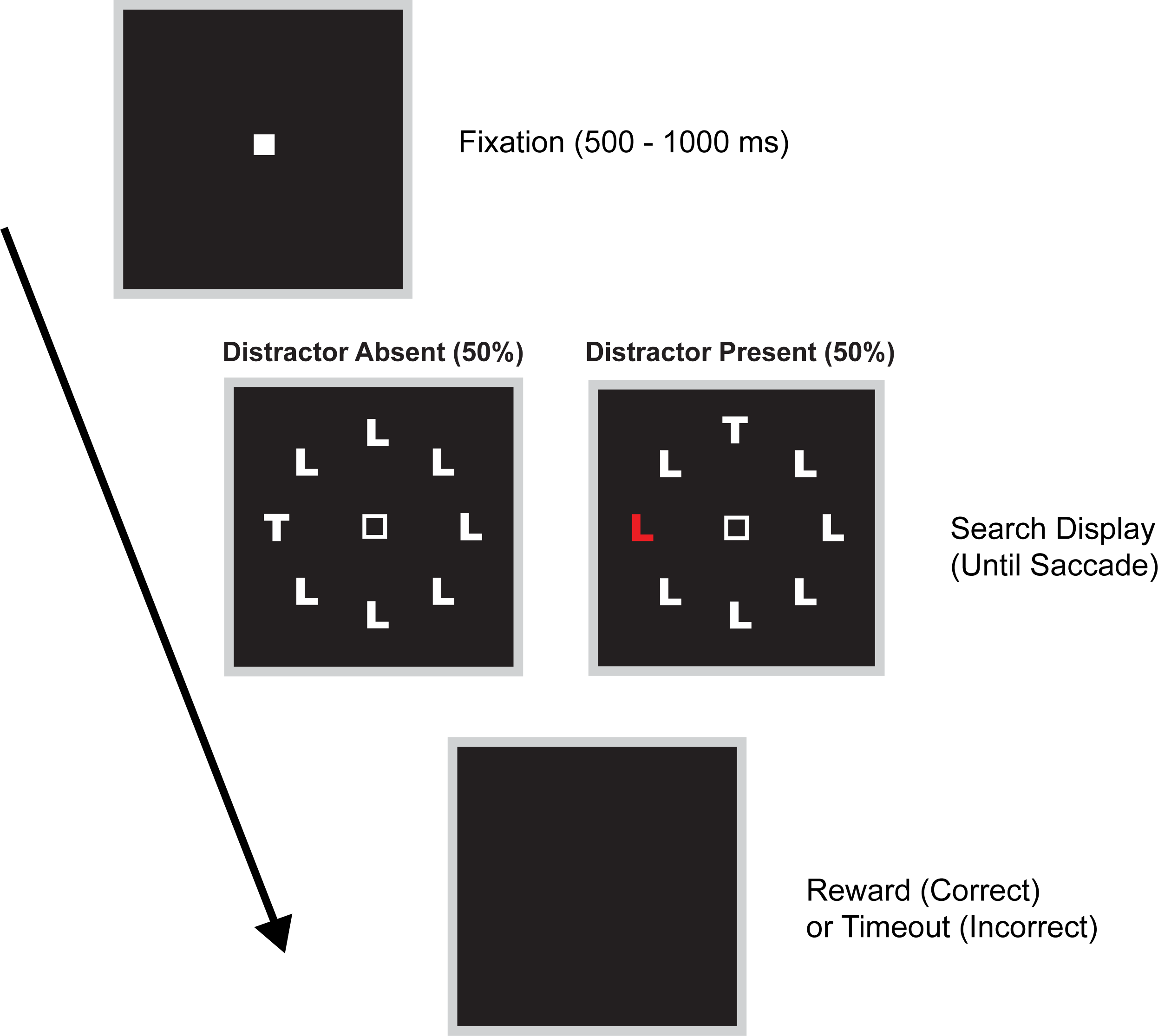
Visual search task. Monkeys fixated for a variable duration (500-1000 ms), at which point the fixation point extinguished and the search array appeared. Monkeys were trained to covertly search for the target, and were rewarded for a first saccade to the target item. A salient, irrelevant distractor appeared unpredictably on half of all trials, which monkey were trained to ignore.

### Suppression of salient distractors by Frontal Eye Field

FEF has been proposed as a source of attention control, acting as a salience map that integrates information about stimulus properties and task goals to bias attention in favor of potentially relevant information [2,21]. Visually responsive neurons in FEF discriminate between target and distractor items during visual search, showing enhanced processing for attended targets vs. unattended distractors prior to a behavioral response [22–25], even when the target item ‘pops out’ of the display on the basis of its bottom-up salience [26,27]. However, it is unknown how visually responsive FEF neurons respond to salient but irrelevant items during visual search. Thus, we contrasted neural responses to task-relevant target items and nonsalient distractors as described in previous work with responses to salient, irrelevant distractors. If salient distractors automatically draw attention as predicted by stimulus-driven hypotheses of distraction control, FEF responses to salient distractor items should be enhanced relative to nonsalient distractor items, paralleling the enhancement observed during selection of task relevant targets.

Replicating previous work, we observed robust selection of task-relevant targets by FEF neurons, with enhanced firing to target items relative to nonsalient distractors (Figure 2B). Critically, we observed no enhancement of responses to salient distractor items. Instead, we observed suppression of the responses relative to both target and nonsalient distractor items (Figure 2B) despite the fact that prefrontal neurons typically show enhanced responses to salient stimuli when they are task-relevant [26,27]. This suggests that the selection of salient stimuli by prefrontal neurons is not automatic, instead depending critically on the relevance of the salient stimulus to ongoing task demands. To examine the latency of these suppression effects, we used running millisecond Wilcoxon rank-sum tests to determine when neural responses to a search target or salient distractor significantly differed from responses to a non-salient distractor when these items fell within or outside of the preferred receptive field (p < 0.01 for 10 consecutive ms). This analysis revealed enhancement of the target 90 ± 15 ms after the onset of the search display, and suppression of the salient distractor at 86 ± 17 ms.

**Figure 2.**
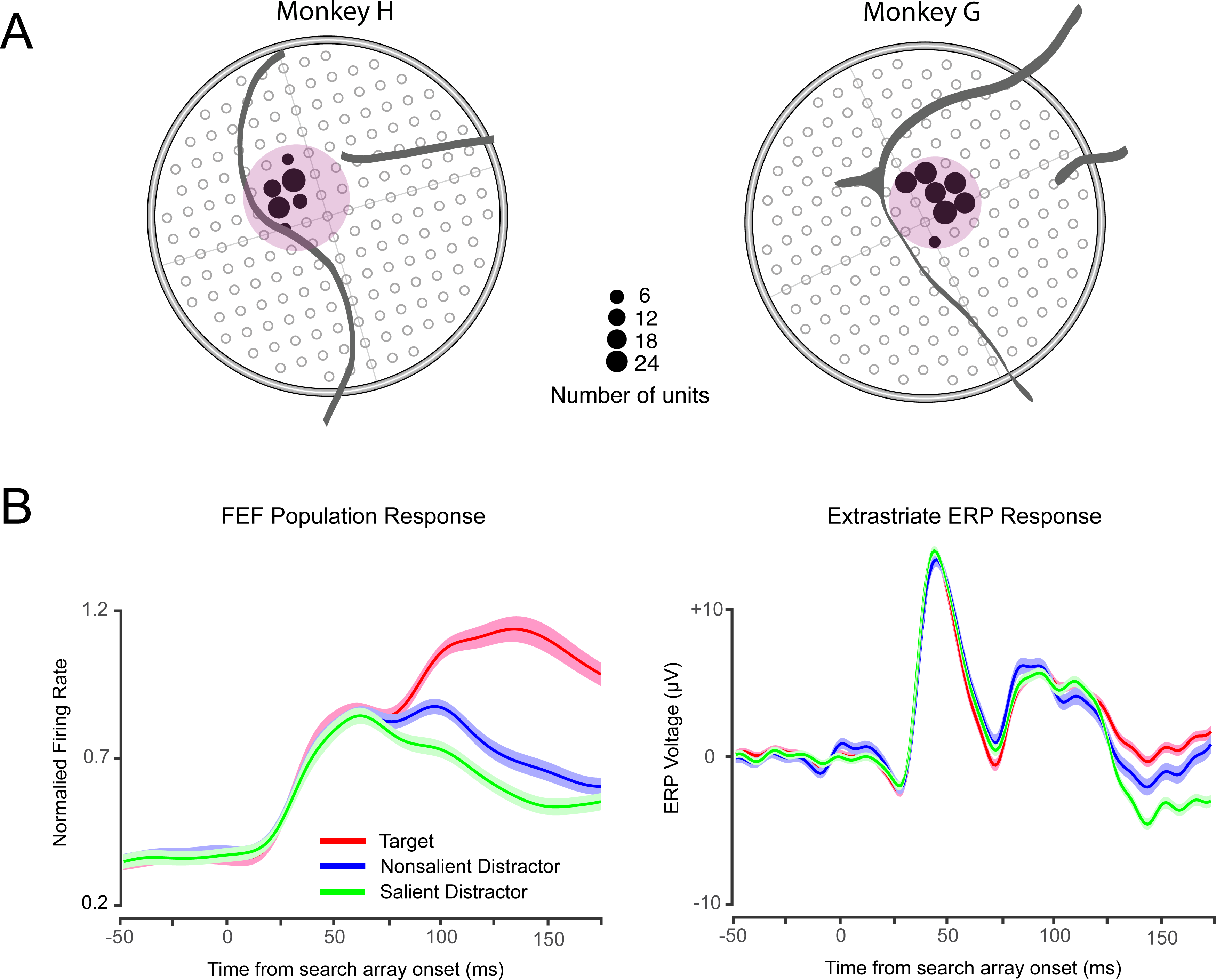
(A) Penetration maps (top) for recordings in each monkey including the total number of units isolated at each location, regardless of task-related modulation. (B) The left panel shows the mean ± SEM of population responses of FEF neurons when the target, nonsalient distractor, or salient distractor fell within a neuron’s receptive field, and the right panel shows the mean ± SEM extrastriate ERP response at electrode site OR to target, nonsalient distractor, or salient distractor appearing in the contralateral (left) hemifield. Data are time-locked to presentation of the search array, and responses were truncated 10ms prior to the saccade.

As noted above, the stimulus-driven capture hypothesis proposes that attention is first drawn to the distractor item and then redirected to the task-relevant target. Even though we observed no evidence of salient distractor selection in FEF firing rates, it is plausible that the presence of selection at sites upstream from FEF may incur a cost in the timing of neural target selection in FEF neurons. Therefore, we also compared when the target was discriminated from nonsalient distractors in the presence and absence of the salient distractor appearing outside of the receptive field. The stimulus-driven capture hypothesis predicts that target selection will be delayed in the presence of the salient distractor. However, we observed no influence of the salient distractor on the latency of target selection by FEF neurons (Distractor Present = 90 ± 18 ms, Distractor Absent = 88 ± 13 ms, *t*(78)=1.20, p=0.23). The absence of salient distractor effects on monkeys’ behavioral performance and FEF responses contradicts the stimulus-driven hypotheses but is consistent with the signal suppression hypothesis. This suggests that the representation of a salient distractor item is proactively suppressed before it can influence neural selection processes and subsequent behavior.

### Target selection and distractor suppression in FEF is carried out by overlapping neuronal populations

To determine whether target enhancement and distractor suppression in FEF are implemented by functionally overlapping or segregated populations of neurons, we examined the firing characteristics of individual attention-related FEF neurons in response to targets and salient distractors. Of the 119 units with significant visual responses, 79 (66%) showed significant target selection, 51 (42%) showed salient distractor suppression, and none (0%) showed significant salient distractor enhancement (Figure 3A). Both target selection and distractor suppression were a consistent feature of visually responsive FEF cells (Figure 3B; also see figure S1). Indeed, of the 51 neurons showing significant salient distractor suppression all also showed target enhancement (Figure 3A). Thus, there appears to be a subclass of neurons within FEF that participate in both processes, providing evidence that only neurons that encode information about task-relevant targets participate the suppression of salient distractors. This finding underscores the benefit of the resolution provided by direct neuronal recordings, as scalp electrophysiological studies could not have provided such an insight into the neural mechanisms of distraction control.

**Figure 3.**
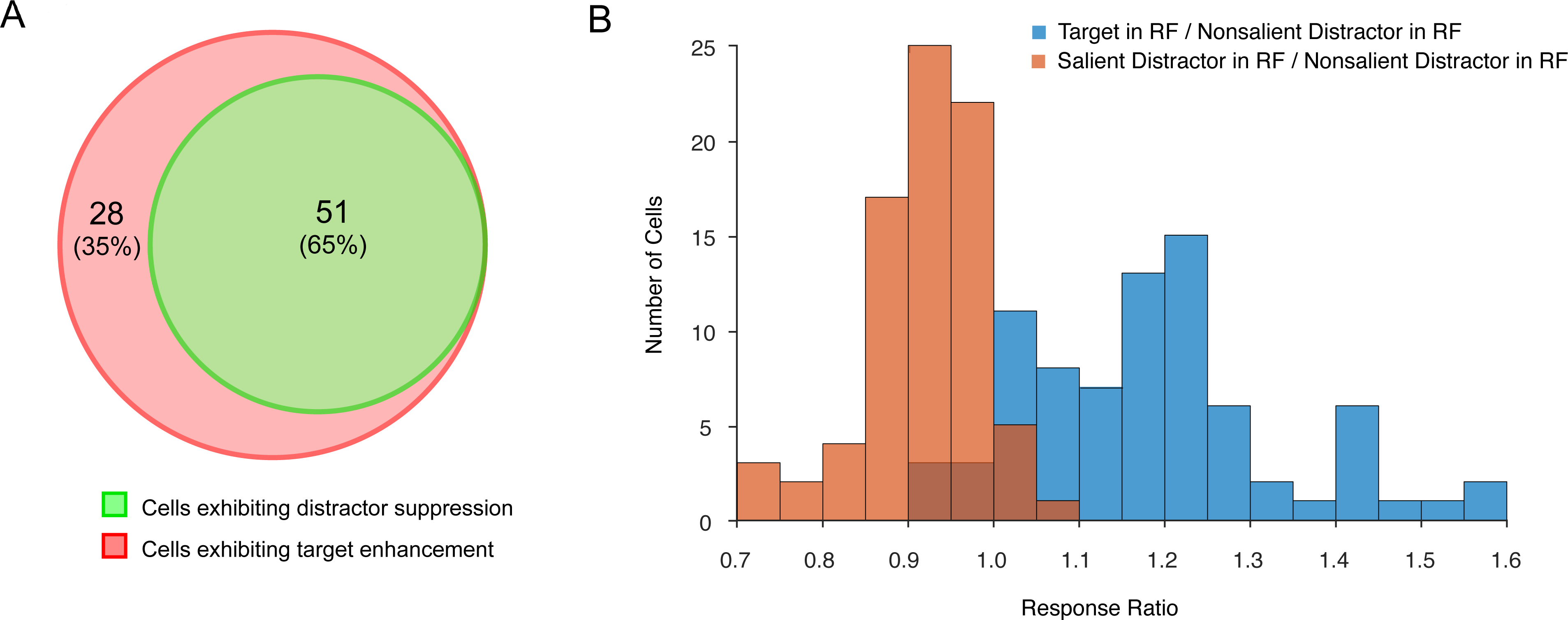
(A) Venn diagram displaying the number of neurons showing target enhancement, distractor suppression, or both. No cells showed distractor suppression in isolation, with all cells showing suppression of the salient distractor also showing enhancement of the target. (B) Distribution of target selection and distractor suppression for all cells showing a significant visual response. The response ratio was calculated by dividing the magnitude of responses targets or salient distractors by responses to nonsalient distractors in the 50-150ms following presentation of the search display. Values greater than one indicate enhancement of responses and values less than one indicate suppression of responses. Both target enhancement and distractor suppression were consistent features in the population of FEF responses.

### Extrastriate ERP responses to salient distractors

Human ERP studies using a task identical to that employed here have played a central role in the debate over the neural mechanisms of distraction control during visual search tasks, providing covert measures of attention in the absence of behavioral distraction. These studies have taken advantage of two lateralized ERP components conjectured to index the selection (*N2pc*) and suppression (*Pd*) of visual information in extrastriate cortex [15,28,29]. A primary goal of the current work was to provide evidence that a putative suppression-related ERP signal, the Pd, reflects the operation of neural suppression processes implemented by FEF. By placing the salient distractor item at a position contralateral to the recording electrode and placing the target on the vertical midline, or vice versa, we isolated the lateralized responses to the different items at extrastriate electrode sites [13,15,30]. To determine the relationship between FEF neuronal responses and extrastriate ERPs, we analyzed trials in which the hemifield contralateral to our extrastriate electrode (electrode site OR) overlapped the receptive field of the population of FEF neurons included in our firing rate analyses, allowing us to directly compare responses across areas and recording modalities (see supplemental experimental procedures). We compared ERP responses to target or salient distractor items with those of the nonsalient distractor item when each appeared in the hemifield contralateral to the recording electrode.

When examining ERP responses to the target in the presence of a salient distractor item, we observed a positive deflection in the ERP response (Figure 2, top panel) relative to the nonsalient distractor occurring 135 ± 25 ms after the search array appeared. This corresponds to the monkey homolog of the human *N2pc* [31–33], and indicates selection of the task-relevant target. The latency of this response was 45 ms later than the target selection signal observed in FEF single neurons (Figure 4). This delay is consistent with previous work suggesting prefrontal cortex as the source of attentional modulation in both extrastriate single-unit and ERP responses [11,27,32,33].

**Figure 4.**
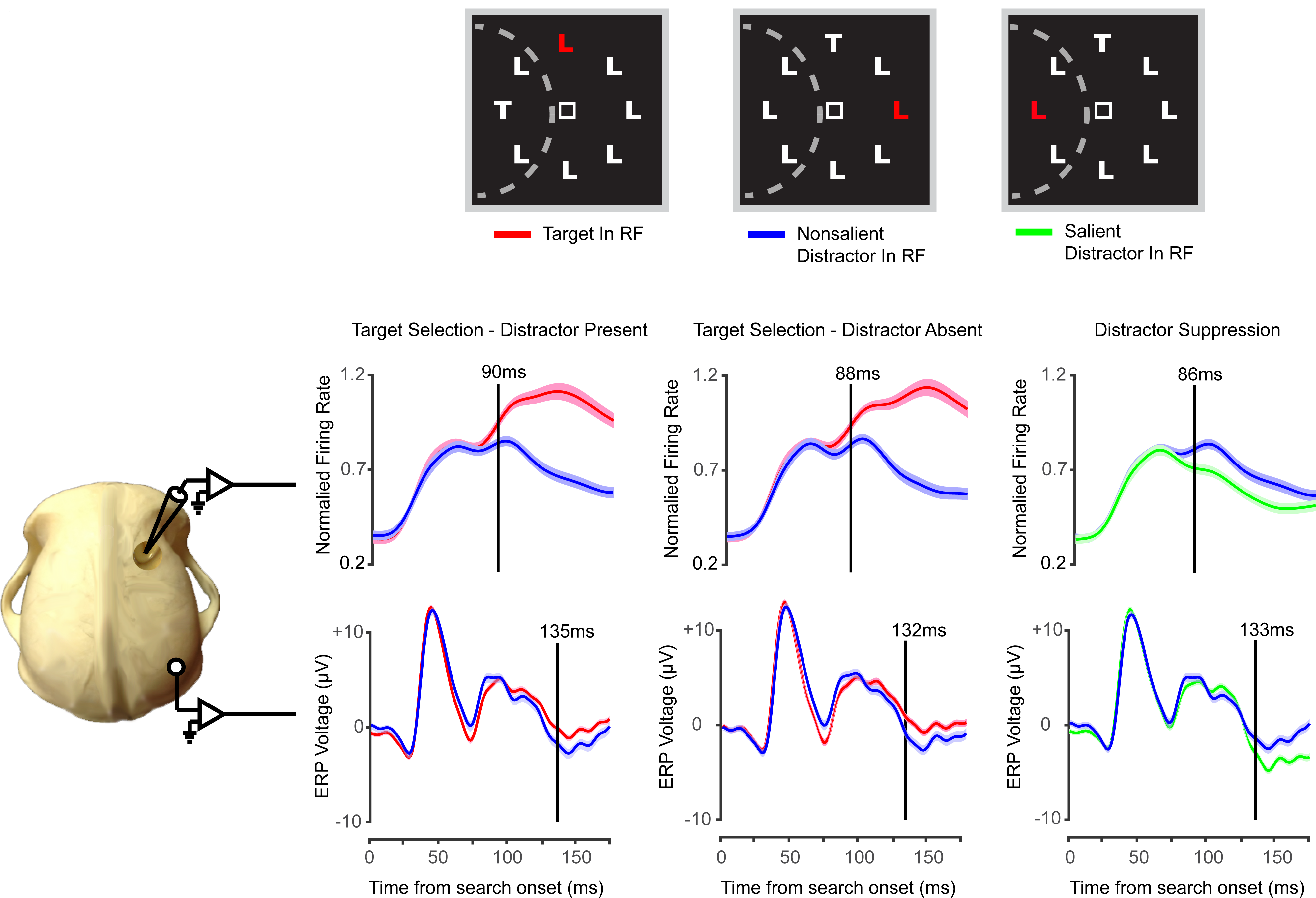
Layout of stimulus displays in each condition, with dotted line corresponding to hypothetical receptive field of FEF neurons in the left visual hemifield. Plots represent target selection (red vs. blue lines) and distractor suppression (green vs. blue lines) in FEF neuronal responses (top panel) and extrastriate ERPs (bottom panel). Vertical lines indicate the point at which responses significantly deviated from one another using a Wilcoxon test (p < .01 for 10 consecutive milliseconds). As can be seen in the figures, both target selection and distractor suppression emerged first in FEF followed by extrastriate ERP responses.

Next, we examined ERP responses to the salient distractor and observed a negative deflection in the ERP response relative to the nonsalient distractor (Figure 2), with this negativity occurring 133 ± 21 ms after search onset. This negativity appears to be the monkey homologue of the *Pd* component observed in the human ERP signal under identical conditions, which is proposed to index attentional suppression processes [13,15,30]. Critically, this distractor suppression effect emerged 47 ms after the suppression observed in FEF neurons (Figure 4), suggesting that, like the *N2pc*, this component reflects the operation of attention control processes driven by prefrontal cortex [32,33]. This observation provides the first evidence linking a putative attentional-suppression related ERP component with suppression of neuronal firing in prefrontal cortex, establishing the scalp-recorded Pd component as a noninvasive readout of prefrontal attentional suppression processes.

It is worth noting that in humans, the *N2pc* and *Pd* components occur with a similar time course and scalp distribution, but are opposite in polarity – the *N2pc* manifests as a negativity at electrode sites contralateral to task-relevant targets whereas the *Pd* appears as a positivity contralateral to a salient distractor. We have previously demonstrated that the monkey homologue of the *N2pc* is inverted in polarity relative to humans (appearing as a positivity in monkeys), which we believe is due to differences in the cortical gyral and sulcal morphology of extrastriate cortex across species [31,32]. The results above show that responses to the distractor, indexing a monkey homologue of the *Pd*, also show a polarity inversion relative to humans (appearing as a negativity in our monkeys). This suggests the possibility that both components reflect a common anatomical source, and may provide a direct readout of attentional modulation in extrastriate cortex across species [29].

## Discussion

Our data are the first to demonstrate FEF contributions to distractor suppression, complementing its well-described role in target selection. This corroborates previous work showing similar target selection and distractor suppression in parietal cortex [34], providing further evidence for a mechanistic overlap in the systems responsible for these processes. We also provide the first demonstration tying this suppression to a nonhuman primate ERP signal of distractor suppression, indicating a homology in extrastriate ERP markers of attentional suppression processes across humans and nonhuman primates. The finding that target selection and salient distractor suppression in FEF neurons preceded ERP responses related to these processes supports previous assertions that FEF is responsible for modulating processing in extrastriate visual areas [3,5,6,32,33]. Taken together, our results are consistent with signal suppression hypotheses that propose an active suppression of distracting information before it can capture attention [11,12]. Thus, when distraction control is successful, the same prefrontal-extrastriate circuit responsible for enhancing task-relevant visual information also participates in the suppression of task-relevant information. Consequently, electrophysiological markers of attention suppression in humans may reflect the effectiveness of distractor control processes implemented by prefrontal cortex, providing a tool for understanding prefrontal control over distraction in both the healthy and disordered brain.

## Acknowledgments

This work was supported by National Institutes of Health Grants F32-EY03922, T32-EY07135, R01-EY019882, R01-EY08890, P30-EY008126, and Robin and Richard Patton through the E. Bronson Ingram Chair in Neuroscience. Requests for materials should be addressed to J.D.C or J.D.S. (, <u>)</u>.

## Notes

**Conflicts of interest:** None

